# A motor endurance assay for zebrafish larvae: proof of principle in chemical and genetic models of neurological disease

**DOI:** 10.64898/2026.10.07.757328

**Authors:** Nuno A. S. Oliveira, Leonor Pereira Lima, Marta Duque, Brígida R. Pinho, José Bessa, Jorge M. A. Oliveira

**Affiliations:** UCIBIO Applied Molecular Biosciences Unit, Mitochondria and Neurobiology Lab, Faculdade de Farmácia, Universidade do Porto, Porto, 4050-313, Portugal; i4HB Institute for Health and Bioeconomy, Universidade do Porto, Porto, 4050-313, Portugal; PhD CAML Doctoral Programme of the Lisbon Academic Medical Centre, Faculdade de Medicina, Universidade de Lisboa, Lisbon, 1649-028, Portugal; IBMC Instituto de Biologia Molecular e Celular, Universidade do Porto, Porto, 4200-135, Portugal; i3S Instituto de Investigação e Inovação em Saúde, Universidade do Porto, Porto, 4200-135, Portugal

**Keywords:** Zebrafish, behaviour, motor endurance, fatigability, amyotrophic lateral sclerosis, TDP-43

## Abstract

Motor fatigability is an early feature of neurological disease, including amyotrophic lateral sclerosis (ALS), but motor endurance assays for zebrafish are largely restricted to juvenile or adult animals. Here, we developed a gravity-fed counter-current swimming assay to quantify motor endurance in 5–7 days post-fertilisation (dpf) zebrafish larvae, which supports the parallel testing of multiple larvae. We detected an increase in motor endurance induced by exposure to caffeine, and a reduction induced by the dopaminergic toxin 1-methyl-4-phenylpyridinium (MPP^+^). In these tests, motor endurance showed a greater dynamic range than spontaneous locomotion. Further, we generated and tested transgenic zebrafish larvae with neuron-restricted overexpression of human TDP43 (hTDP43), the major pathological protein in ALS patients. Zebrafish with neuronal hTDP43 expression showed ALS-relevant hTDP43 mislocalisation and significantly reduced motor endurance at 5 dpf. This method for motor endurance assessment in 5–7 dpf zebrafish is simple, sensitive, and enables the characterisation of motor fatigability in models of human neurological disease (TDP43-linked ALS and MPP^+^-induced parkinsonism).

## Introduction

Zebrafish (*Danio rerio*), especially in the larval development stages, have notable advantages as biomedical models, regarding both their experimental tractability and their genetic and physiological similarity to humans. The small size, optical transparency, and rapid development of larval zebrafish allow the screening of large numbers of individuals, enabling rapid phenotypic characterisation in a vertebrate system (Patton et al., 2021).

Fatigability and loss of motor endurance are hallmarks of neurological impairment or reduced fitness in humans and model animals (Brooks and Dunnett, 2009; Hannan et al., 2025; Kluger et al., 2013; Rouviere et al., 2012). Specifically, focal muscle weakness and fatigability are early findings in amyotrophic lateral sclerosis (ALS) (Ravits et al., 2026), a neurodegenerative disease defined by motor neuron loss, with TDP43 neuropathology present in >95% of cases (Hardiman et al., 2017). In the context of ALS, motor endurance assessment in rodent models (e.g. using a rotarod treadmill) has proven useful for advancing disease understanding and identifying therapeutic targets (Van Hoecke et al., 2012; Zheng et al., 2026).

Considering the high tractability, throughput, and growing availability of larval zebrafish models, motor endurance assessment in these models can enable further mechanistic and target validation studies. However, existing motor endurance assays were developed for adult zebrafish. These assays consist of counter-current swimming tasks in purpose-made swimming tunnels, and have been used to enrich thecharacterisation of zebrafish models of ALS (Hannan et al., 2025; Ramesh et al., 2010). These counter-current assays for adult zebrafish generate turbulent and fast water streams that larval zebrafish cannot overcome. In parallel, established locomotor tracking methods for zebrafish larvae mostly focus on spontaneous locomotion and reflex responses (Fero et al., 2011) — to our knowledge, as far as we could find in the literature, no standardised method for motor endurance assessment in sub-14 days post-fertilisation zebrafish larvae has been reported. Thus, the lack of a standardised motor endurance assay for zebrafish larvae constitutes a significant gap in the characterisation of these models.

This gap represents a current limitation of larval zebrafish studies, compared with adult fish and rodents, for which standardised tests (e.g. swimming tunnels for fish, the rotarod for mice) are well established (Brooks and Dunnett, 2009; Ramesh et al., 2010). Here we bridge the gap between larval locomotor analysis and adult endurance assessment by developing a new gravity-fed counter-current swimming assay to assess motor endurance in zebrafish larvae. We leverage the genetic tractability and high-throughput potential of zebrafish to validate this new motor endurance assay. Furthermore, we provide proof of principle that this assay can detect concentration-dependent effects of pharmacological modulators, and a disease-relevant motor phenotype in zebrafish models of parkinsonism and ALS.

## Methods

### Motor Endurance Assay

Motor endurance in zebrafish larvae was assessed with the gravity-fed counter-current swimming system developed in the present study (Fig. 1A, B). This system consists of two reservoirs placed at different heights – a lower reservoir (*R*_1_) feeding the swimming tunnel (ST) with a water-level-dependent water flow (*F*_1_); and an upper reservoir (*R*_2_) feeding the lower reservoir with flow rate *F*_2_. The difference in the water levels of the two reservoirs (*h*_2_) is significantly larger than the water level in the lower reservoir (*h*_1_), and was adjusted such that the inflow to the lower reservoir (*F*_2_) is always higher than the outflow (*F*_1_). This causes the water level in the lower reservoir (*h*_1_) to increase gradually over time, leading to a corresponding increase in flow rate at the swimming tunnel (*F*_1_) (Fig. 1D, E).

**Fig. 1.**
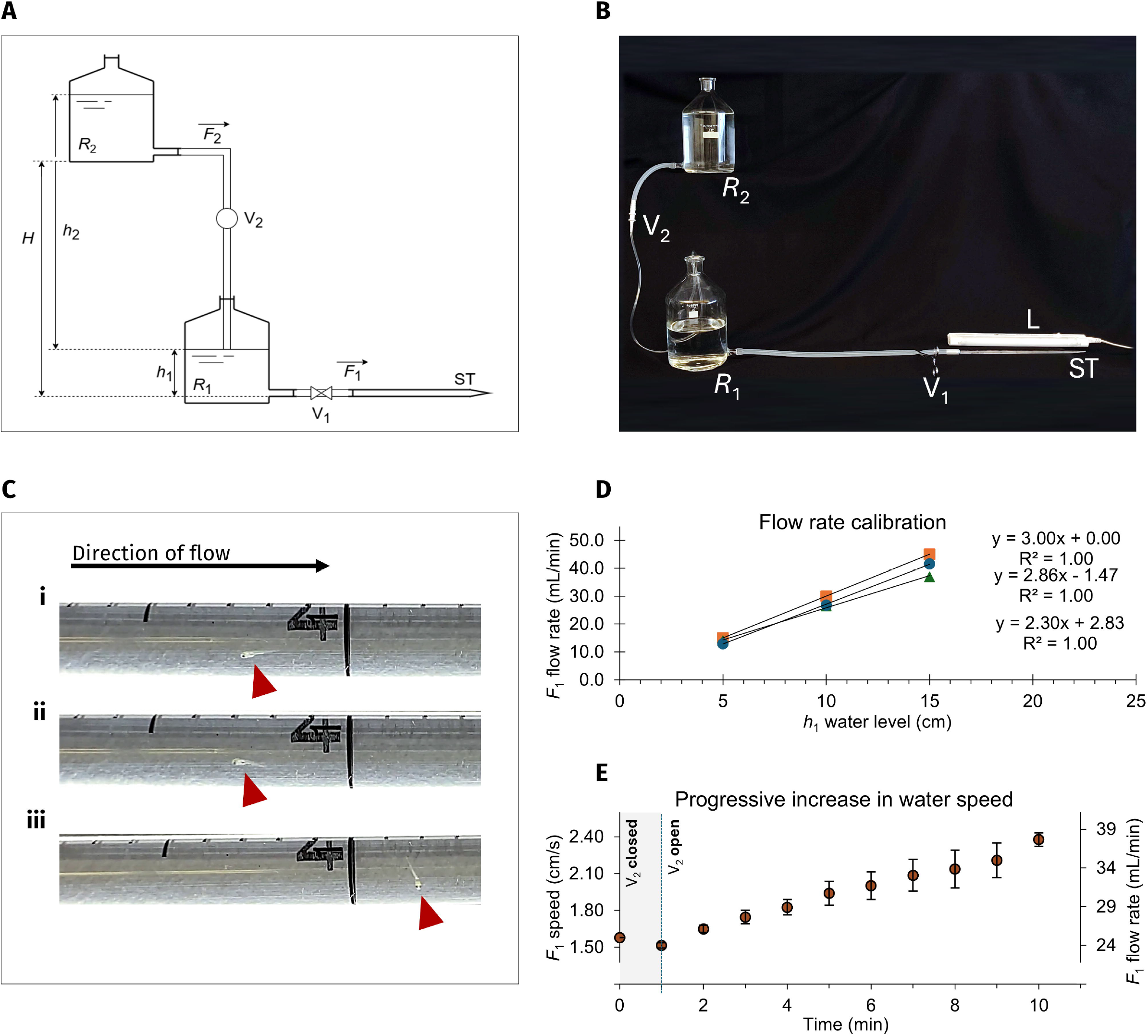
Gravity-fed counter-current swimming assay for motor endurance. **(A)** Schematic representation of the setup used for the motor endurance assay. Water flow rate (*F*_1_) in the swimming tunnel (ST) is proportional to the water level (*h*_1_) in the lower reservoir (*R*_1_) and controlled by a clamp (V_1_). A second reservoir (*R*_2_), placed 42 cm (*H*) higher than *R*_1_, feeds the lower reservoir with a flow rate *F*_2_, controlled by a screw valve (V_2_). *F*_2_ is proportional to the difference in water levels of *R*_2_ and *R*_1_ (*h*_2_). The height difference between the two reservoirs ensures that *F*_2_ (inflow to the lower reservoir) exceeds *F*_1_ (outflow from the lower reservoir). This causes *h*_1_, and therefore *F*_1_, to increase over time, progressively increasing water speed in the ST and, consequently, the required swimming effort. **(B)** Photograph of the setup, with a fluorescent lamp (L) illuminating the swimming tunnel. **(C)** Representative images of a 5 dpf zebrafish larva at different stages during the motor endurance assay: aligned with the flow (i), swimming against the flow (ii), and being carried downstream by the flow (iii). **(D)** Flow rate is linearly proportional to water level *h*_1_; calibration curves obtained from three pipettes used as swimming tunnels. Based on these calibration curves, the initial water level *h*_1_ is adjusted so that the initial flow rate is 25 mL/min (water speed of 1.58 cm/s). **(E)** Flow rate in the swimming tunnel (*F*_1_) and water speed (*v*_F1_) increase progressively over time, starting at 1 minute, when V_2_ is opened, causing *h*_1_ to rise. Data are mean ± SEM from three pipettes used as swimming tunnels.

We implemented the assay using two 2 L Pyrex aspirator bottles at a 42 cm height difference (*H*, Fig. 1A, 1B). The bottles were connected by a plastic tube (4 mm inner diameter) with a plastic screw valve for flow regulation. A pliable silicone tube (8 mm inner diameter) was used to connect the lower bottle to a 5 mL plugged serological pipette (Sarstedt; 5.8 mm inner diameter), which served as “the swimming tunnel”. Laboratory film (Parafilm “M”) was used to ensure a water-tight seal between the swimming tunnel and the silicone tubing. Water temperature was kept at 26 ± 1 ºC using 10 W USB-powered miniature water heating elements inside the reservoirs. Flow calibration was performed by measuring the flow rate at three different water levels *h*_1_, using a graduated cylinder (Fig. 1D). Initial flow rate was then set to 25 mL/min by adjusting the initial water level in *R*_1_. This initial flow rate corresponds to a water speed of 1.6 cm/s inside the swimming tunnel (Eqns. S2).

For motor endurance assessment, 5–7 days post-fertilisation (dpf) zebrafish larvae were first moved to a 90 mm dish with zebrafish embryo medium, 15 minutes prior to the experiment, for acclimation. 5 to 10 larvae were then moved into the swimming tunnel, through the tip, with a Pasteur pipette. After a 5-minute adaptation period and ensuring that the larvae had moved beyond the mid-point of the pipette, water flow in the swimming tunnel was initiated by opening the V_1_ clamp. One minute later, the upper reservoir valve (V_2_) was opened, initiating the water flow *F*_2_. The flow rate at the swimming tunnel (*F*_1_) then increases gradually over time (Fig. 1E), up to a maximum rate of 38 mL/min (water speed of 2.4 cm/s) after 10 minutes. Endurance times were recorded as the time interval from the start of the assay until each larva failed to maintain its position in the swimming tunnel and was ejected by the water flow (Fig. 1C). Since zebrafish larvae show thigmotaxis, with a preference to touch the walls instead of being in the centre of a space (Bagatto et al., 2002), the tube was manually rotated 180º once per second throughout the assay, preventing larvae from adhering to the pipette walls.

### Free swimming recording

The video recordings and analyses were performed similarly to how we previously described (Silva et al., 2022), using 96-well plates for motor tracking. 5 dpf morphologically normal larvae with demonstrated swimming ability were placed in the wells and, after 15 min of acclimation, their swimming activity was videorecorded for 20 min, at 27 ± 1 °C, with a high-resolution digital camera (Basler ace acA1440-220um). The videos were acquired at 25 frames per second, and the positions of zebrafish larvae were extracted in real time, using a custom MATLAB script (Code available: https://github.com/nuno-SO/Oliveira2026_zebrafishTracking). The tracking data were analysed using MATLAB and Megabouts (Jouary et al., 2025), to extract the swimming trajectory and total distance swum per larva.

### Zebrafish Husbandry

Adult AB-strain zebrafish (*Danio rerio*) were supplied by the Interdisciplinary Centre of Marine and Environmental Research (CIIMAR, Portugal). Tg(*elavl3*:*GAL4FF*^ccu5Tg^); Tg(*UAS*:*mCherry*) double transgenic zebrafish were obtained from the Orger Lab at Champalimaud Research (Champalimaud Foundation, Portugal). Zebrafish were maintained at 28 ± 1 °C, under 14 h–10 h light-dark cycles and eggs were produced as we previously described (Oliveira et al., 2024). Briefly, on the afternoon before egg collection, 1–2 zebrafish couples were placed in 1.1 L breeding tanks. On the following morning, 1 h after the start of the light phase, the eggs were collected, washed and maintained in E3 medium (5.0 mM NaCl, 0.17 mM KCl, 0.33 mM CaCl_2_, 0.40 mM MgCl_2_ (Cold Spring Harbor Laboratory Press, 2011)) at 28 °C. Handling of zebrafish followed the European Directive 2010/63/EU and the Portuguese Law (Decreto-Lei 113/2013). The animal facility (BOA-ICBAS) was licensed by *Direção-Geral de Alimentação e Veterinária (DGAV)* and the experiments were approved by the ICBAS-UP Animal Welfare Body (P564/2025/ORBEA-ICBAS).

### Zebrafish drug exposures

Exposure to MPP^+^ was achieved by immersing 3-dpf wild-type zebrafish in 500 µM MPP^+^ iodide (Sigma D048) diluted in E3 medium, for 48 hours, in 24-well plates (5 fish/well/500 μL) in a semi-static system (Silva et al., 2022), with media changes every 24 hours. Acute effects of caffeine were studied by exposing 5-dpf wild-type zebrafish to freshly-prepared 50–300 μM anhydrous caffeine (Sigma W222402), diluted in E3 medium, for 15 minutes (Wilson et al., 2024). Stock solutions of MPP^+^ and caffeine were prepared in ultrapure water (solvent control).

### Generation of transgenesis vectors

Vectors for zebrafish transgenesis were assembled using the Gateway recombination-based Tol2Kit (Kwan et al., 2007). Human *TARDBP* cDNA (from Addgene #27462,Johnson et al., 2009) was restriction-cloned to the entry vector pME-MCS. The resulting vector was then recombined with p5E-*UAS* (containing the inducible *UAS* element) and p3E-polyA (containing the SV40 late polyadenylation signal sequence) into the destination vector pDestTol2CG2 (which includes a *cmlc2*:EGFP transgenesis marker) using Gateway LR Clonase II Enzyme mix (ThermoFisher), according to the manufacturer’s protocol.

Standard chemical transformation was performed with NZY5α competent cells (NZYTech), grown overnight at 37□°C. Vector selection was performed with 100□μg/ml Ampicillin (Normon) in the growth medium. Plasmids were purified with NZYMiniprep kit (NZYTech) and confirmed by Sanger sequencing. Final plasmids were purified with phenol/chloroform and concentration was determined by NanoDrop 1000 Spectrophotometer (ThermoFisher).

### In vitro mRNA synthesis, microinjection and transgenesis

Transgenic Tg(*UAS:TARDBP*; *cmlc2*:*EGFP*) zebrafish were generated through TOL2-mediated transgenesis (Kwan et al., 2007). *Tol2* cDNA was transcribed by Sp6 RNA polymerase (ThermoFisher) after Tol2-pCS2FA (Addgene #133032) vector linearization with NotI restriction enzyme (ThermoFisher). *Tol2* mRNA was purified as previously described (Bessa et al., 2009). One-cell stage embryos were injected with 1 nL solution containing 25□ng/µL of transposase mRNA, 25□ng/µL of purified plasmid, and 0.05% phenol red (Sigma-Aldrich). Microinjected embryos were selected and grown until adulthood. Transgenic F0 fish were outcrossed with wild-type adults, and their progeny were screened using *cmlc2:EGFP* fluorescence as a surrogate for transgene insertion.

Transgenic Tg(*elavl3*:*Gal4FF*) zebrafish (ccu5Tg) were generated by co-injecting *elavl3*:*Gal4FF* plasmid (Chen et al., 2013, 12 ng/ul) with a *UAS* reporter plasmid for a green fluorescent protein (CaMPARI2) and Tol2 transposase mRNA (50 ng/ul) into 1–2 cell stage eggs from a cross of Tübingen with *mitfa*^-/-^ fish. Embryos showing a green signal at 3–4 dpf were raised and screened as adults for germ line transmission. The founder for the Gal4FF driver line was selected from progeny of positive animals that showed a broad and stable expression pattern, and crossed with a Tg(*UAS*:*mCherry*) reporter line.

Neuron-restricted expression of human TDP43 was achieved by crossing zebrafish from the Tg(*UAS:TARDBP*; *cmlc2*:*EGFP*) line with zebrafish from theTg(*elavl3*:*GAL4FF*); Tg(*UAS*:*mCherry*) (ccu5Tg) driver line (pan-neuronal GAL4). The resulting progeny was then selected based on EGFP and mCherry fluorescence, to identify fish expressing TDP43 and fish expressing GAL4 only (spawn-matched controls).

### Western blot

Protein extraction and quantification were performed as we previously described (Oliveira et al., 2024). 5–15 zebrafish larvae (5 dpf) per condition were washed and transferred to ice-cold RIPA lysis buffer (50 mM Tris-HCl, pH 7.4, 1% NP-40, 0.25% sodium deoxycholate, 1 mM EDTA, 0.1% SDS; 10 μL per larva) supplemented with protease inhibitor cocktail (Thermo Scientific). After a freeze-thaw cycle and mechanical homogenisation (Precellys Evolution, 6800 rpm, 5 × 10 s with 15 s pauses, CK28 beads), lysates were centrifuged at 2,000×g (10 min, 4 °C) and supernatant protein concentrations were quantified by the Bradford method.

After total protein quantification, samples were denatured (95 °C, 5 min) in Bolt LDS sample buffer (Invitrogen) supplemented with Bolt sample reducing agent (Novex), loaded (20 μg protein) into 4–12% Bolt Bis-Tris Mini Protein Gels (ThermoFisher), electrophoresed (200 V, 22 min), and transferred into PVDF membranes (30 V, overnight at 4 °C). Membranes were then blocked (1 h, room temperature) with 5% bovine serum albumin (BSA, NZYTech) in PBS with 0.05% Tween-20 (PBST), and incubated with anti-TDP43 rabbit monoclonal antibody (Cell Signaling D9R3L, 1:1000). After washing with PBST, membranes were incubated for 1 h with goat anti-rabbit secondary antibody conjugated with horseradish peroxidase (G-21234, 1:4000, Thermo Scientific), for detection using Novex ECL Chemiluminescent kit (Invitrogen) and a ChemiDoc MP Imaging System (Bio-Rad). Membrane Coomassie (Coomassie Brilliant Blue R-250, ThermoFisher) staining was used as loading control.

### Wholemount immunofluorescence

7 dpf zebrafish larvae samples were processed for immunofluorescence as we previously described (Oliveira et al., 2024). Briefly, after fixation with 4% paraformaldehyde, larvae were permeabilised with PBS with 1% Triton X-100 (PBTr) and dehydrated with methanol. After rehydration, the samples were blocked, incubated with anti-TDP43 rabbit monoclonal antibody (Cell Signaling D9R3L, 1:200, 16 h);washed with PBTr, and incubated with the respective Alexa-488-conjugated secondary antibody (A-11034, Life Technologies; 1:200, 16 h). They were then washed with PBS, treated with 1 μg/mL Hoechst 34580 for 30 minutes for nuclear counter-staining, and mounted in agarose topped with 80% glycerol. Imaging was conducted with the previously described Nikon-Prior-Hamamatsu Imaging System (Pinho et al., 2019), equipped with a 20× objective (Nikon CFI Plan Fluor 20X, NA0.5) and a DAPI/FITC/TRITC/CY5 Quad band-pass filter (Chroma 89401). Excitation wavelengths were 405 nm for Hoechst 34580 (emission collected between 425 and 460 nm) and 488 nm for Alexa-488 (emission collected between 500 and 545 nm).

### Data analysis and statistics

Numerical data are primarily shown as mean ± standard error of the mean (SEM) of the *n* number indicated in figure legends, or as boxplots with median, mean, quartiles, range (most extreme data points that fall within 1.5 times the IQR from the quartiles), and outliers (Tukey, 1977). Numbers of independent spawns used for zebrafish experiments are stated in figure legends. Normality assumptions were tested with the Shapiro-Wilk test; square root or logarithm transformations were used for non-normal data before statistical testing through factorial ANOVA with Tukey’s post-hoc, using a linear model that considered the treatment conditions, with independent spawning events as a blocking factor (i.e. effect ∼ condition + spawn). *p*-values under 0.05 were considered statistically significant. Data analyses were performed with R 4.3.0 (R Core Team, 2021) in Rstudio 2023.06.0, using the *tidyverse* (Wickham et al., 2019), *rstatix* (Kassambara, 2023a), *ggpubr* (Kassambara, 2023b), *drc* (Ritz et al., 2015) and *cowplot* packages (Wilke, 2020).

## Results

### Development of a motor endurance assay for zebrafish larvae: a gravity-fed counter-current assay achieves a predictable and laminar flow regime

Motorised counter-current swimming tests have been used to assess motor endurance in adult zebrafish, but they create a fast turbulent flow that is not suitable for young (5-7 dpf) zebrafish larvae (Hannan et al., 2025; Ramesh et al., 2010). Thus, to assess motor endurance in zebrafish larvae, we developed a new counter-current swimming assay. We aimed to create a predictable, laminar water flow, strong enough to elicit counter-current swimming in zebrafish larvae, while allowing easy observation of their performance, without requiring mechanized pumps. We assembled a new gravity-fed system, inspired by previously described systems used for counter-current training of zebrafish larvae during development (Bagatto et al., 2002; Fiaz et al., 2012), rather than for motor endurance assessment.

To achieve a swimming tunnel with progressively-accelerating laminar flow, we assembled a gravity-fed system with two reservoirs at different heights (Fig. 1A, B). Water flows from the upper reservoir to the lower reservoir, increasing the water level in the lower reservoir. Consequently, the water flow rate (mL/min) from the lower reservoir to the swimming tunnel, and its internal water speed (cm/s), progressively increases over time (Fig. 1D-E; see Methods for more details). This setup remains in a laminar flow regime (Reynolds number 105 < *Re* < 165, well within the conventional range of Re < 2300 for laminar flow (Reynolds, 1883; White, 2008); Eqn. S3), which induces the counter-current swimming reflex in zebrafish larvae without immediately overpowering them (Fig. 1C).

For our swimming tunnel dimensions (5.8 mm inner diameter) and a typical 5-dpf zebrafish body length of 4.2 mm (Müller et al., 2008), our target flow rates of 25–38 mL/min correspond to water speeds of 3.8–5.7 body lengths per second (BL/s), a common size-normalised measure of fish swimming performance, or 1.6 to 2.4 cm/s (Eqn. S2). These speeds exceed the maximum sustained swim speed of zebrafish at this stage, which is approximately 3 BL/s (Fiaz et al., 2012) — thus ensuring that all larvae eventually reach the assay endpoint (ejection from the swimming tunnel). In summary, our two-reservoir gravity-fed setup induces predictable laminar counter-current flow suitable for motor endurance assessment in 5–7 dpf zebrafish larvae.

### Motor endurance is a viable metric in zebrafish larvae

After validating the water flow conditions in our experimental setup, we characterised the counter-current swimming performance of wild-type, control zebrafish larvae. We observed zebrafish swimming in bouts, alternating between periods of swimming and periods of coasting (Movie S1). Endurance times displayed a right-skewed distribution, with a mean endurance time of 197 s ± 22.5 s (Fig. 2A-i). After a square-root transformation, data were normally distributed and suitable for parametric analysis (Fig. 2A-iii).

**Fig. 2.**
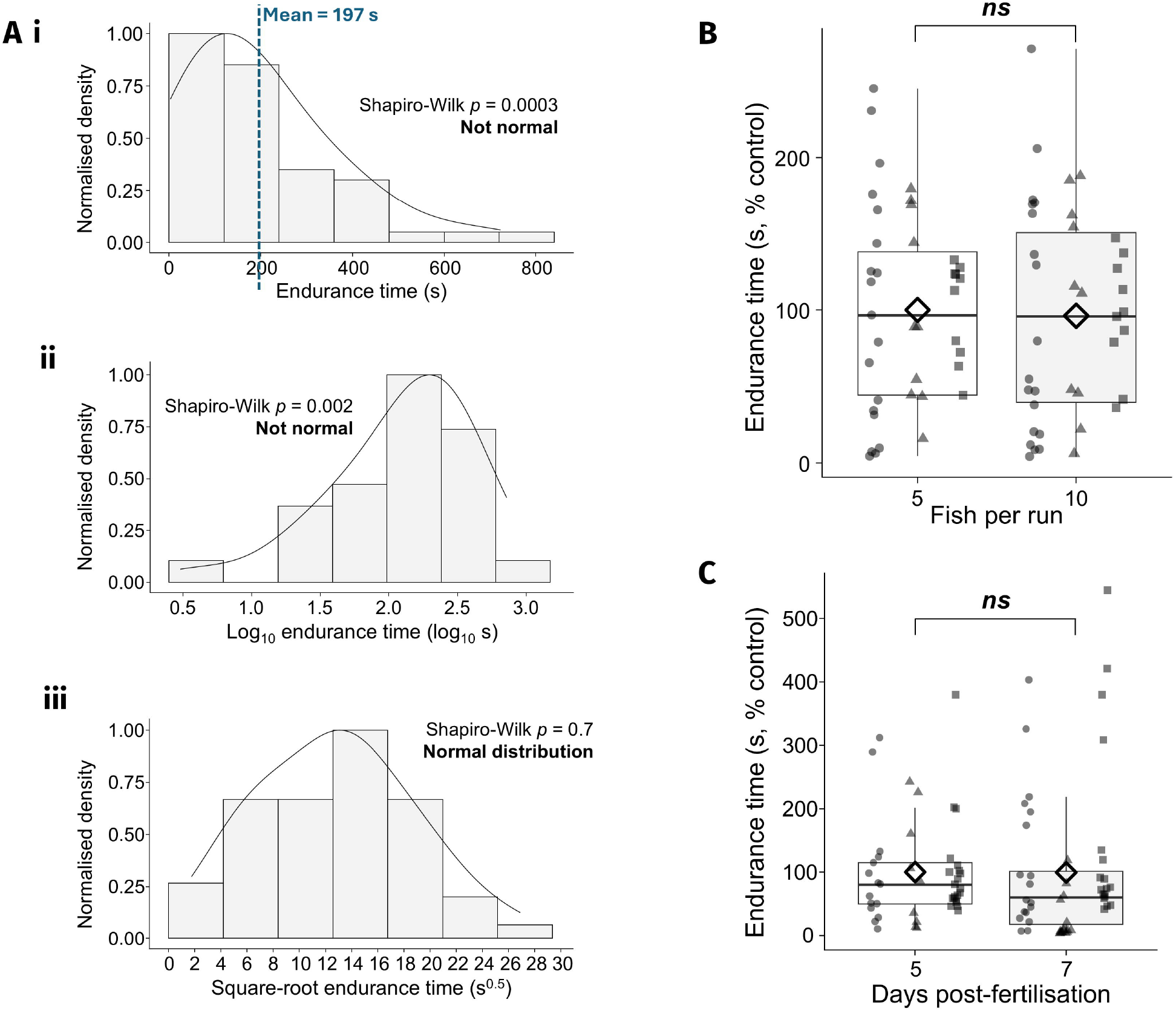
Motor endurance characterisation in control zebrafish larvae. **(A)** Distribution of motor endurance data from 5 dpf wild-type AB zebrafish, shown on linear (i), logarithmic (ii), and square-root (iii) scales. Data are from *n* = 53 zebrafish from 3 independent spawns; histograms show bars and a density line scaled to a maximum of 1. Square-root-transformed endurance times follow an approximately normal distribution (*p* = 0.7, i.e. not significantly different from normal distribution; Shapiro-Wilk normality test). **(B)** Mean endurance time (time until failure in % control) is similar when runs start with 5 or 10 fish inside the tunnel. *n* = 39 fish per condition, at 5 dpf. (**C**) Mean endurance time is similar at the ages of 5 and 7 dpf. *n* = 45–52 fish per condition, with 5 loaded in the tunnel in each run. *ns*, not significant, *p* > 0.05, ANOVA on square-root-transformed data, with spawn included in the ANOVA as a blocking factor. Differently-shaped individual points in boxplots indicate different zebrafish spawns, diamonds indicate global means.

To assess the impact of the number of fish in the swimming tunnel and of fish age, we compared motor endurance results under different conditions. Evaluating 5 or 10 fish per run in the swimming tunnel did not significantly change motor endurance times (Fig. 2B). Performance was also identical between 5 dpf and 7 dpf zebrafish larvae (Fig. 2C).

These results show that time endured in this counter-current swimming assay is a viable metric for the study of motor endurance in zebrafish larvae. In the next sections, we focus on comparing groups of larvae subjected to different treatments, to assess the applicability of this motor endurance assay to biomedical research.

### Motor endurance detects both stimulation and inhibition by pharmacological modulators

To evaluate the effect of different pharmacological treatments on motor endurance, we tested zebrafish larvae exposed to either a stimulant: caffeine (adenosine receptor antagonist); or a movement inhibitor: MPP^+^ (dopaminergic neurotoxin). We selected concentration ranges that have been shown to affect zebrafish behaviour (Pinho et al., 2016; Silva et al., 2022; Wilson et al., 2024). Exposure to caffeine induced a concentration-dependent increase in motor endurance, with a calculated EC_50_ of 111 μM and a maximum increase of approximately 3-fold vs. solvent control (Fig. 3A). We compared this result with a classical spontaneous swimming assay, on a 96-well plate. Caffeine induced an increase in spontaneous swimming distance, with a stable plateau effect between 50 and 150 μM (∼1.5 fold increase) (Fig. 3B). Thus, for caffeine exposure, the motor endurance assay showed higher dynamic range (progressive increase between 50 and 150 µM) and reached maximal plateau effect at a higher concentration (300 µM), compared with spontaneous swimming (50 µM). We then chose a single-point concentration to assess the statistical significance of the caffeine effect (130 μM for motor endurance and 50 μM for spontaneous swimming) and observed a statistically significant increase in both motor endurance and spontaneous swimming (Fig. 3C). Lastly, to assess if a drug known to induce hypoactivity would reduce motor endurance time, we tested the effect of the parkinsonian toxin MPP^+^ at 500 μM (Silva et al., 2022). As expected, MPP^+^ induced a reduction in both motor endurance and spontaneous swimming (Fig. 3D), validating the bi-directional responsiveness of the motor endurance assay.

**Fig. 3.**
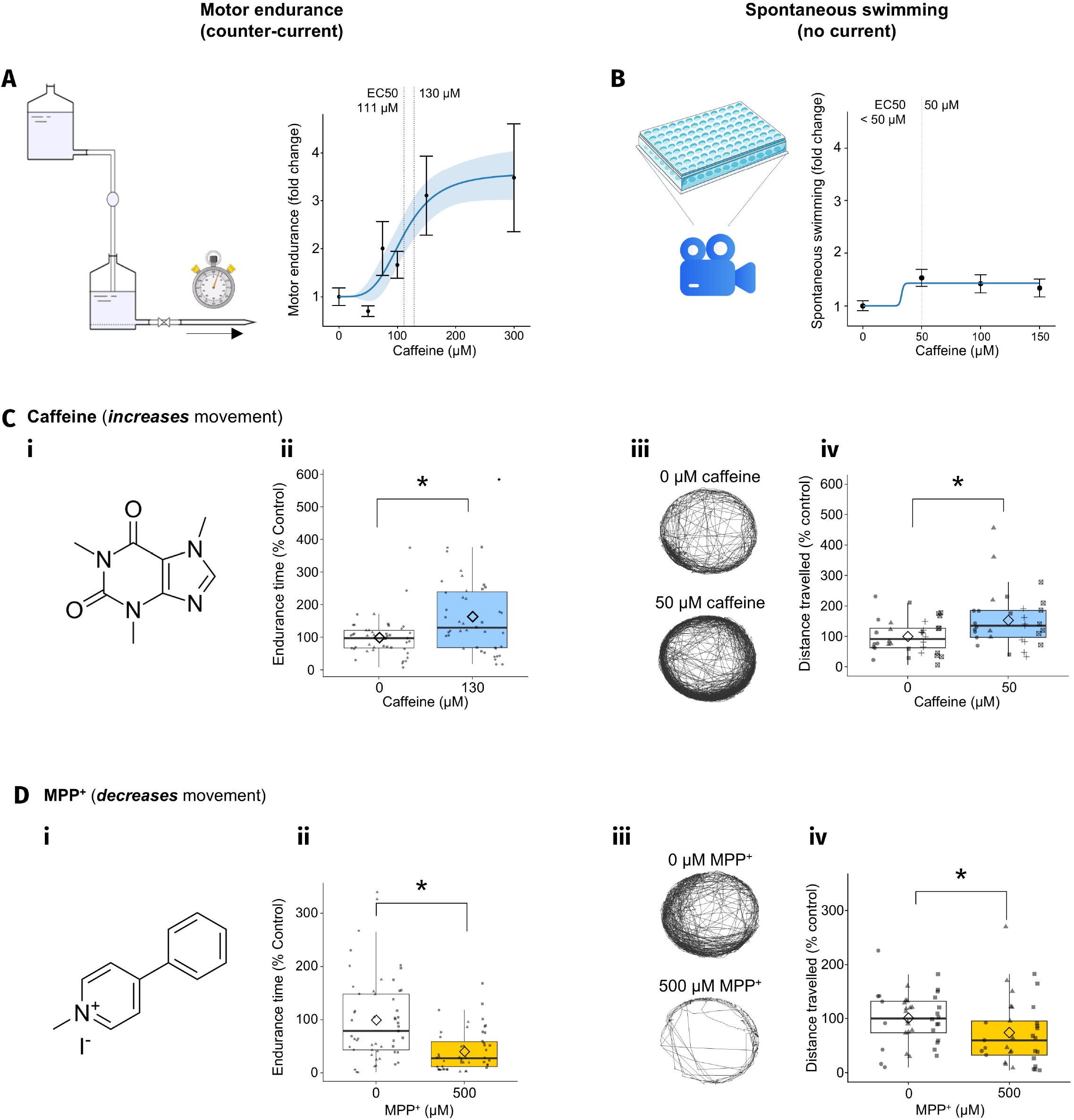
The motor endurance assay detects stimulation by caffeine and inhibition by the dopaminergic toxin MPP^+^. **(A)** Fold change in motor endurance of 5 dpf zebrafish larvae exposed to different caffeine concentrations, ranging from 50 to 300 μM, relative to solvent control. Data are mean ± SEM with a fitted log-logistic dose-response curve (fixed lower asymptote). Dashed vertical lines mark the EC_50_ (111 ? 21 μM) and the concentration used in single-point experiments (130 μM). **(B)** Fold change in spontaneous swimming activity (total distance swum over 20 minutes) of 5 dpf zebrafish larvae exposed to different caffeine concentrations, ranging from 50 to 150 μM, relative to solvent control. Data are mean ± SEM with a fitted log-logistic dose-response curve (fixed lower asymptote). Dashed vertical line marks the concentration used in single-point experiments (50 μM). **(C)** Caffeine exposure increases both motor endurance and spontaneous swimming. (i) Molecular structure of caffeine; (ii) motor endurance time of 5 dpf zebrafish exposed to130 μM caffeine, vs. solvent control; (iii) representative spontaneous swimming trajectories of 5 dpf zebrafish exposed to 50 μM caffeine, vs. solvent control; (iv) spontaneous swimming distance of 5 dpf zebrafish exposed to 50 μM caffeine, vs. solvent control. * *p* < 0.05, ANOVA on log-transformed data, with spawn included as a blocking factor, followed by Tukey’s multiple comparisons; *n* = 31–45 zebrafish per condition from 4 independent spawns. **(D)** MPP^+^ exposure decreases both motor endurance and spontaneous swimming. (i) Molecular structure of MPP^+^; (ii) motor endurance time of 5 dpf zebrafish exposed to 500 μM MPP^+^, vs. solvent control; (iii) representative spontaneous swimming trajectories of 5 dpf zebrafish exposed to 500 μM MPP^+^, vs. solvent control; (iv) spontaneous swimming distance of 5 dpf zebrafish exposed to 500 μM MPP^+^, vs. solvent control. * *p* < 0.05, ANOVA on square-root-transformed data, with spawns included in the ANOVA as a blocking factor, followed by Tukey’s multiple comparisons; *n* = 33–53 zebrafish per condition, from 3–4 independent spawns. Differently-shaped individual points in boxplots indicate different zebrafish spawns, diamonds indicate means.

### ALS-model zebrafish show reduced motor endurance

Following the assay with the chemical model MPP^+^, we next assessed the applicability of the motor endurance assay to a genetic model of neurodegeneration. We generated ALS-model fish, induced by overexpressing human TDP43 in zebrafish neurons. To achieve the TDP43 overexpression, we used the GAL4/UAS system. This binary system induces high expression levels of the gene of interest (TDP43), under the control of the *UAS* element, dependent on the presence of the GAL4 transcriptional activator (Fig. 4A). Thus, by crossing neuronal-GAL4 transgenic zebrafish, Tg(*elavl3*:*GAL4FF*), with conditional-TDP43 zebrafish, Tg(*UAS*:*TDP43*), we can induce neuronal TDP43 expression. We validated TDP43 expression by western blot (Fig. 4B) and whole-mount immunofluorescence (Fig. 4D), confirming that human TDP43 can be successfully expressed in the neurons of zebrafish larvae and shows ALS-like cytoplasmic mislocalisation.

**Fig. 4.**
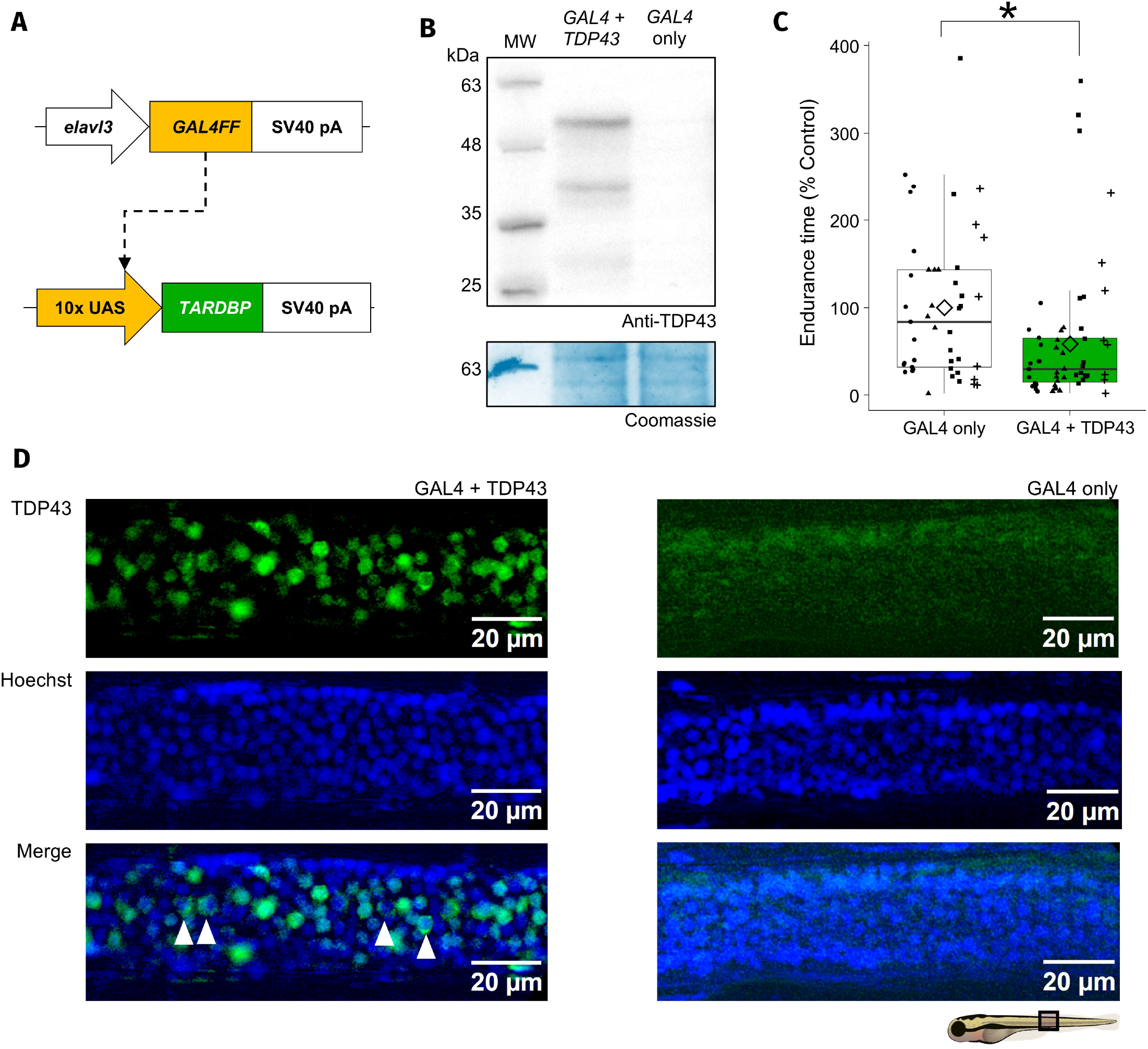
ALS-model fish show reduced motor endurance. **(A)** Construct used for pan-neuronal overexpression of human TDP43. An *elavl3* promoter drives neuronal expression of the transcriptional activator GAL4FF, which strongly activates the *UAS* element controlling TDP43 expression. **(B)** The Tg(*elavl3*:*GAL4FF*); Tg(*UAS*:*TDP43*) double-transgenic construct successfully induces TDP43 expression, as observed in an anti-TDP43 western blot of protein extracts from 5-dpf double-transgenic zebrafish (*GAL4 + TDP43*), compared with single-transgenic Tg(*elavl3*:*GAL4FF*) control fish from the same spawn (*GAL4*). Both full-length TDP43 and its smaller C-terminal fragments are visible. **(C)** Motor endurance times of 5 dpf zebrafish with pan-neuronal TDP43 overexpression, vs. spawn-matched controls; *p* < 0.05, ANOVA on log-transformed data; *n* = 45–55 zebrafish per condition from 4 independent spawns. Differently-shaped individual points in boxplots indicate different zebrafish spawns, diamonds indicate means. **(D)** Representative anti-TDP43 immunofluorescence (green) labelling of the spinal cords of 7-dpf zebrafish with pan-neuronal TDP43 overexpression, vs. spawn-matched controls. Nuclei are stained with Hoechst 34580 (blue). Note the cytoplasmic mislocalisation of TDP43 in some neurons (arrowheads).

Then, we tested the motor endurance of zebrafish overexpressing human TDP43, vs. controls from the same spawn. We observed a statistically significant reduction in the motor endurance of zebrafish expressing human TDP43 (Fig. 4C), supporting an ALS-relevant impairment of motor endurance.

## Discussion

Increased fatigability and reduced endurance are common findings in human health conditions (Kluger et al., 2013). Larval zebrafish offer tractability and throughput advantages as disease models for mechanistic and drug screening studies (Bashirzade et al., 2022); however, current motor endurance tests for zebrafish are restricted to older fish (Hannan et al., 2025; Ramesh et al., 2010). Here we present a motor endurance assay for early zebrafish larvae (as young as 5 dpf), show representative data, describe the effect of drug exposures on motor endurance, and apply this new assay to a new ALS-relevant zebrafish model of neuronal TDP43 proteinopathy.

Our motor endurance assay is inspired by previously described counter-current swimming setups, used to study the effects of swim training on zebrafish larvae (Bagatto et al., 2002; Fiaz et al., 2012). However, swim training is conducted at water speeds below the critical swimming speed (maximum prolonged swimming velocity; Cano-Barbacil et al., 2020). Counter-current swimming below the critical swimming speed has also been used to investigate the mechanisms underlying rheotaxis in zebrafish, showing that both visual and mechanosensory input contribute to this behaviour, and that zebrafish can use flow velocity gradients as navigational cues (Newton et al., 2023; Oteiza et al., 2017). In contrast, our assay uses flow speeds above the critical speed to induce high-intensity swimming and relatively rapid fatigue. This high-intensity, rapid-fatiguing swimming regime, testing up to 10 larvae at a time, increases the throughput of the assay, compared with critical speed determination studies. Critical speed studies provide more detailed data but have lower throughput, due to the requirement of long swimming intervals (several minutes) at each flow speed, for each individual (Ramesh et al., 2010). Additionally, the use of pre-feeding-stage zebrafish larvae eliminates differences in dietary intake as a source of variability, and greatly reduces body length variability, compared with studies in older zebrafish.

We also build on a previously described open-source system that uses a microcontroller-based flume to evaluate swimming performance in zebrafish starting at 14 dpf (Widrick et al., 2018). This microcontroller-based system allows finer control over flow specifications, including the ability to generate precise and sustained flow velocities—these characteristics make it ideal for critical swim speed determination over a range of timepoints. Our system complements the microcontroller-based flume by providing a simpler, non-motorised setup, that can be used to quickly assess the motor endurance of large numbers of early zebrafish larvae (5–7 dpf), and that has been validated with chemical modulators and a genetic disease model.

We observed that the effects of drug treatments on our counter-current motor endurance assay broadly matched their effects on spontaneous locomotion. These results support the hypothesis that evoked swimming intensity and swim bout initiation are correlated, even though they are controlled by at least partially independent mechanisms (Holman et al., 2023). Specifically, here we tested MPP^+^, a dopaminergic toxin known to impair locomotor behaviour in 5 dpf zebrafish (Thirumalai and Cline, 2008), that has been used as a model of parkinsonism (Silva et al., 2022); and caffeine, an adenosine receptor antagonist and stimulant. In our data, MPP^+^ exposure reduced spontaneous locomotion and motor endurance, consistent with its known dopaminergic toxicity (Christensen et al., 2020; Silva et al., 2022). In contrast, exposure to 130 μM caffeine increased motor endurance and spontaneous locomotion, in line with previous data showing that acute caffeine exposure at this concentration increases average swimming speed in adult zebrafish (Wilson et al., 2024). In summary, our new counter-current motor endurance assay was able to detect changes in swimming endurance,which are related to, but separate from changes in spontaneous locomotion. This suggests potential to provide insights into motor circuits and performance components that are not fully captured by free-swimming locomotion tracking alone.

Since we were able to modulate motor endurance with drug exposures, we also characterised the motor endurance of ALS-model zebrafish overexpressing human TDP43. ALS primarily affects motor function; thus, a motor endurance assay may provide a disease-relevant readout that complements spontaneous locomotion assessment. We observed that TDP43 overexpression lowered motor endurance, as expected for a model of ALS (Ramesh et al., 2010), highlighting the potential of motor endurance assessment in larval zebrafish models of disease.

In conclusion, in this study we developed a new method for motor endurance assessment in zebrafish larvae. We showed that performance in this endurance assay is sensitive to interventions affecting motor control, including drug exposures and genetic modelling of neurological diseases. Compared with critical swim speed evaluation, our simple motor endurance assay assesses higher-intensity swimming performance, which increases experimental throughput at the cost of providing a simpler measurement of endurance. We propose combining this new motor endurance assay with spontaneous locomotion and critical swim speed assessment—thus gathering insights on independent components of motor control and performance and broadening the behavioural characterisation of zebrafish models of disease.

## Supporting information

Supplemental Equations S1-S3

## Resource availability

### Lead contact

Requests for further information and resources should be directed to and will be fulfilled by the lead contact, JMAO.

### Materials availability

Plasmids and reagents generated in this study are available upon reasonable request.

## Data and code availability

Data reported in this paper will be shared by the lead contact upon request.

Original code reported in this paper is available on https://github.com/nuno-SO/Oliveira2026_zebrafishTracking.

Any additional information required to reanalyse the data reported in this paper is available from the lead contact upon request.

## Acknowledgements

The Tg(*elavl3*:*GAL4FF*) zebrafish line was created and shared by Sabine Renninger and Ruth Diez del Corral from the Orger Lab (Champalimaud Centre for the Unknown, Portugal).

Funding for JMAO’s laboratory: FCT – Fundação para a Ciência e a Tecnologia, Portugal – (UIDB/04378/2025; UIDP/04378/2025; NORTE2030-FEDER-02707400; 2024.16934.PEX). NASO: “la Caixa” Foundation (ID 100010434) PhD fellowship - LCF/BQ/DR23/12000018. LPL: FCT, 2024.01464.BD. JB’s laboratory: FCT project COMPETE2030-FEDER-00731800-16163 and UID/4293/2025. JB’s FCT contract CEECIND/03482/2018.

## Author contributions

NASO: Conceptualization, Formal analysis, Investigation, Methodology, Software, Visualization, Writing – original draft. LPL: Investigation, Writing – review and editing.

MD: Investigation, Methodology, Writing – review and editing. BRP: Resources, Writing – review and editing. JB: Conceptualization, Funding acquisition, Resources, Supervision, Writing – review and editing. JMAO: Conceptualization, Funding acquisition, Project administration, Resources, Supervision, Writing – review and editing.

