## Supplemental Equations S1-S3 for "A motor endurance assay for zebrafish larvae: proof of principle in chemical and genetic models of neurological disease"

**Water speed and Reynolds number calculations**

**S1. System characteristics**

- Tube inner diameter, *d*

$$d=0.58\text{ cm}=0.0058\text{ m}$$

- Tube cross-sectional area, *A*

$$A=\pi\cdot r^{2}=\pi\cdot\frac{d^{2}}{4}=2.64\cdot{10}^{-5}\text{ m}^{2},$$

where *r* is the tube inner radius.

- Volumetric flow rate, *Q*_F1_ (conversion to SI units)

$$Q_{F1, \min}=25 mL min^{-1}=\frac{25\cdot{10}^{-6} m^{3}}{60 s}=4.17\cdot{10}^{-7} m^{3}s^{-1}$$

$$Q_{F1,\max}=38 mL min^{-1}=\frac{38\cdot{10}^{-6} m^{3}}{60 s}=6.33\cdot{10}^{-7} m^{3}s^{-1}$$

- Water properties at 26 °C:
  - Density: $\rho=997\text{ }\text{kg m}^{-3}$
  - Dynamic viscosity: $\mu=8.7 \cdot{10}^{-4} Pa s$

**S2. Flow speed**

- Water velocity in a tube, *v*

$$v=\frac{Q}{A}$$

where *Q* is the fluid flow rate and *A* is the tube cross-sectional area.

$$v_{\text{min}}=\frac{4.17\cdot{10}^{-7}}{2.64\cdot{10}^{-5}}=1.58\cdot{10}^{-2} \text{m s}^{-1} = 1.58 \mathrm{cm}\text{ s}^{-1}$$

$$v_{\text{max}}=\frac{6.33\cdot{10}^{-7}}{2.64\cdot{10}^{-5}}=2.40\cdot{10}^{-2} \text{m s}^{-1} = 2.40 \mathrm{cm}\text{ s}^{-1}$$

- Speed relative to body length of a 5 dpf zebrafish (BL)

$$\text{BL}=4.2\text{ mm}=0.42\text{ cm}$$

$$v_{\text{min, BL}}=\frac{1.58\text{ cm s}^{-1}}{0.42 \text{cm}\text{ }{BL}^{-1}}=3.8 BL s^{-1}$$

$$v_{\text{max, BL}}=\frac{2.40\text{ cm s}^{-1}}{0.42 \text{cm}\text{ }{BL}^{-1}}=5.7 BL s^{-1}$$

**S3. Reynolds number**

- Reynolds number (calculated using SI units)

$$\text{Re}=\frac{\rho\cdot d}{\mu}\cdot v$$

where *ρ* is the fluid density, *d* is the characteristic dimension (inner diameter of the tube), *μ* is the fluid dynamic viscosity, and *v* is the fluid velocity.

$$\text{Re}_{\text{min}}=\frac{997\cdot0.0058}{8.7\cdot{10}^{-4}}\cdot1.58\cdot{10}^{-2}=105$$

$$\text{Re}_{\text{max}}=\frac{997\cdot0.0058}{8.7\cdot{10}^{-4}}\cdot2.40\cdot{10}^{-2}=165$$
